# Single-cell Analysis of Paired FFPE Placentas Reveals Trophoblast Reprogramming and Immune Dysregulation in Chronic Villitis of Unknown Etiology

**DOI:** 10.64898/2026.02.28.708431

**Authors:** Juan Li, Meiling Wang, Quantao Zhang, Tian Luo, Yusheng Chen, Guangjun Yin, Yuanzhen Suo, Yongcheng Wang, Yue Wang, Fengchun Gao

**Author notes:** Corresponding author. (F.G.); wangyue@sdfmu. edu.cn (Y.W.); (YC.W.); (Y.S.). These authors contributed equally to this work (Juan Li, Meiling Wang, Quantao Zhang).

## Abstract

Villitis of unknown etiology (VUE) is a common placental inflammatory lesion associated with adverse pregnancy outcomes, yet its mechanisms remain unclear. Mechanistic studies are hindered by reliance on archived formalin-fixed paraffin-embedded (FFPE) tissues, which are incompatible with standard single-cell methods. Here, we develop an FFPE-compatible single-nucleus RNA sequencing strategy and apply it to paired placentas from sequential pregnancies. VUE placentas exhibit pronounced immune expansion and disrupted trophoblast differentiation. A distinct extravillous trophoblast population and a subset of syncytiotrophoblasts aberrantly express classical MHC class I molecules, promoting natural killer cell engagement and indicating loss of trophoblast immune privilege. Pro-inflammatory maternal macrophages infiltrate, while fetal Hofbauer cells are reduced and transcriptionally reprogrammed. Increased intracellular viral burden and JAK–STAT signaling correlate with aberrant MHC class I expression, suggesting stress-induced local alloimmune activation. These findings define a cellular basis for disrupted maternal–fetal immune tolerance in VUE and demonstrate the utility of FFPE-based single-cell transcriptomics in placental disease.

## INTRODUCTION

Villitis of unknown etiology (VUE) is a common and insidious chronic inflammatory lesion of placenta, histopathologically defined by the infiltration of maternal CD8^+^ T cells into the villous stroma(*1*). Affecting approximately 5%-15% in term pregnancies(*2*) with a recurrence rate of 10%-30%(*3*), VUE is associated with a variety of adverse outcomes, including fetal growth restriction, preterm birth, stillbirth, and neonatal neurodevelopmental disorders(*2*, *4*). Despite its prevalence and profound clinical impact, the cellular and molecular mechanisms driving VUE remain elusive, posing a significant barrier to the development of targeted diagnostics and therapeutics.

Accumulating evidence supports the concept that VUE reflects a breakdown of maternal–fetal immune tolerance(*5*, *6*). This dysregulation involves the aberrant upregulation of MHC class I and class II molecules on placental cells, facilitating the presentation of fetal antigens to the maternal immune system and triggering a targeted alloimmune attack(*7*). While T cells have traditionally been regarded as the primary effectors, recent attention has shifted toward the critical role of macrophages. Quantitative studies have demonstrated that macrophages, rather than T cells, actually constitute the predominant immune population within VUE lesions(*8*). However, the lineage and polarization dynamics of these macrophages remain contentious. While Kim et al. originally suggested they primarily arise from the *in situ* proliferation of resident fetal Hofbauer cells (HBCs)(*9*), subsequent studies have detected a substantial component of infiltrating maternal macrophages(*10*). Furthermore, the polarization state of placental macrophages remains poorly defined, specifically whether the observed shift toward a pro-inflammatory (M1) or anti-inflammatory (M2) phenotype is driven by maternal monocyte recruitment or local HBC reprogramming(*11*). These discrepancies largely stem from the resolution limits of traditional methodologies, such as immunohistochemistry and bulk RNA-sequencing, obscure the complex spatial and functional heterogeneity of the placental microenvironment. These approaches obscure cellular heterogeneity and fail to reliably distinguish maternal from fetal lineages, leaving the precise molecular drivers of VUE incompletely understood.

Single-cell RNA sequencing (scRNA-seq) has transformed the analysis of complex tissues by enabling unbiased, high-resolution characterization of cellular heterogeneity and intercellular communication. However, conventional scRNA-seq requires fresh tissue and high-quality RNA, which are rarely available for VUE, as placentas are routinely fixed and archived after delivery(*12*). Moreover, the rarity of VUE and its substantial inter-individual heterogeneity further complicate prospective sample collection. Recent advances in FFPE-compatible single-cell and single-nucleus RNA sequencing technologies(*13*) now permit transcriptomic profiling of archived specimens, offering a unique opportunity to revisit longstanding mechanistic questions in placental pathology.

In this study, we applied a random-priming based single-nucleus RNA sequencing approach to paired FFPE placental samples collected from sequential pregnancies over a five-year period. This paired design allowed each individual to serve as her own internal control, thereby minimizing confounding effects of genetic background and environmental exposure. Using this strategy, we systematically interrogated the cellular landscape of VUE and identified aberrant trophoblast differentiation and immune activation—centered on MHC class I–expressing trophoblast subsets and macrophage-driven inflammation—as core features of VUE pathogenesis.

## RESULTS

### Paired FFPE placental sampling enables integrated single-cell analysis of VUE with minimal inter-individual heterogeneity

To delineate the cellular basis of villitis of unknown etiology (VUE), we assembled a longitudinal cohort of 12 archived formalin-fixed paraffin-embedded (FFPE) placental specimens obtained from six women across sequential pregnancies (Fig. 1A). Three individuals experienced a normal first pregnancy followed by VUE in the second pregnancy, whereas the remaining three exhibited VUE in both pregnancies. All VUE cases met standardized histopathological criteria for high-grade disease. This paired design allowed each individual to serve as an internal control, thereby minimizing confounding effects of genetic background and environmental exposure.

**Figure 1.**
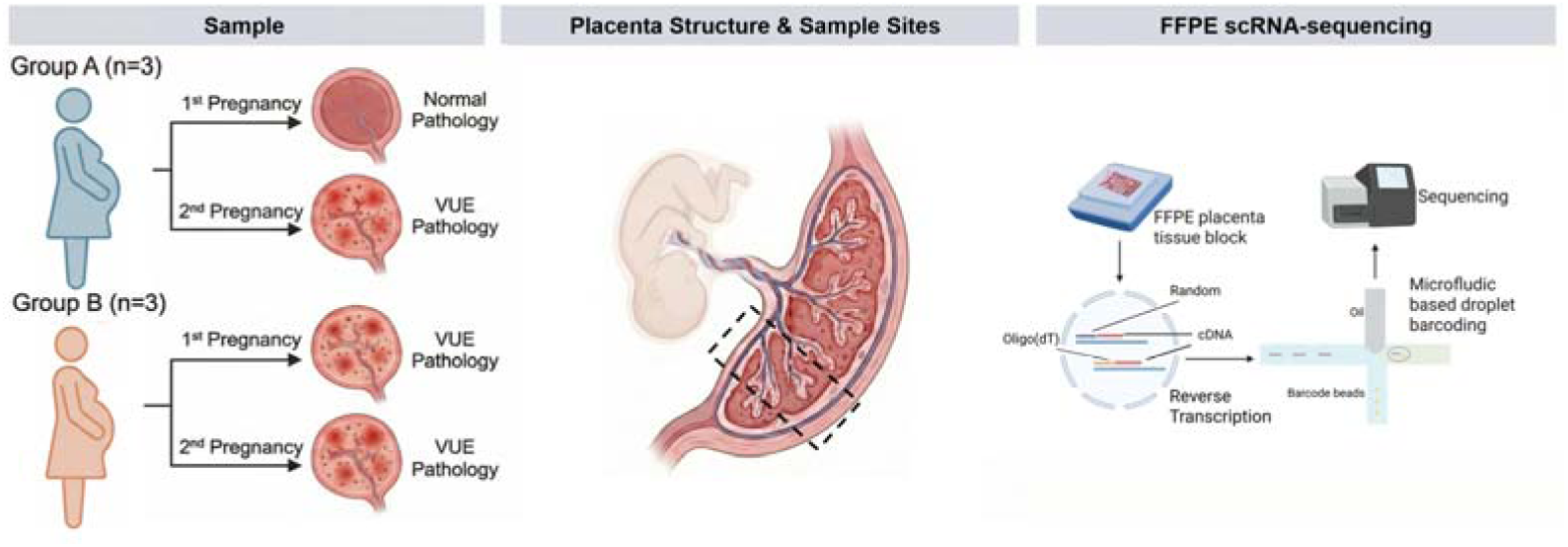
Study design and scRNA-seq workflow. Schematic of the study cohort: Group A (n=3) presented with a normal first pregnancy followed by a VUE-complicated second pregnancy; Group B (n=3) presented with recurrent VUE. Illustration of placental sampling sites (dashed lines). Workflow for single-cell RNA sequencing derived from FFPE tissue blocks.

All samples were processed using a random-priming–based single-nucleus RNA sequencing (snRNA-seq) workflow optimized for FFPE tissues(*14–16*). Following quality control, normalization, and batch correction, the integrated dataset captured the major cellular constituents of the placental microenvironment, including trophoblast lineages, immune populations, and stromal and vascular cells. Uniform manifold approximation and projection (UMAP) visualization demonstrated extensive overlap of cells across individuals and pregnancies, indicating robust dataset integration and minimal batch effects (Fig. 2A, 2B). Unsupervised clustering resolved 20 transcriptionally distinct clusters, which were annotated into 11 major cell types based on canonical marker genes (Fig. 2C–2G).

**Figure 2.**
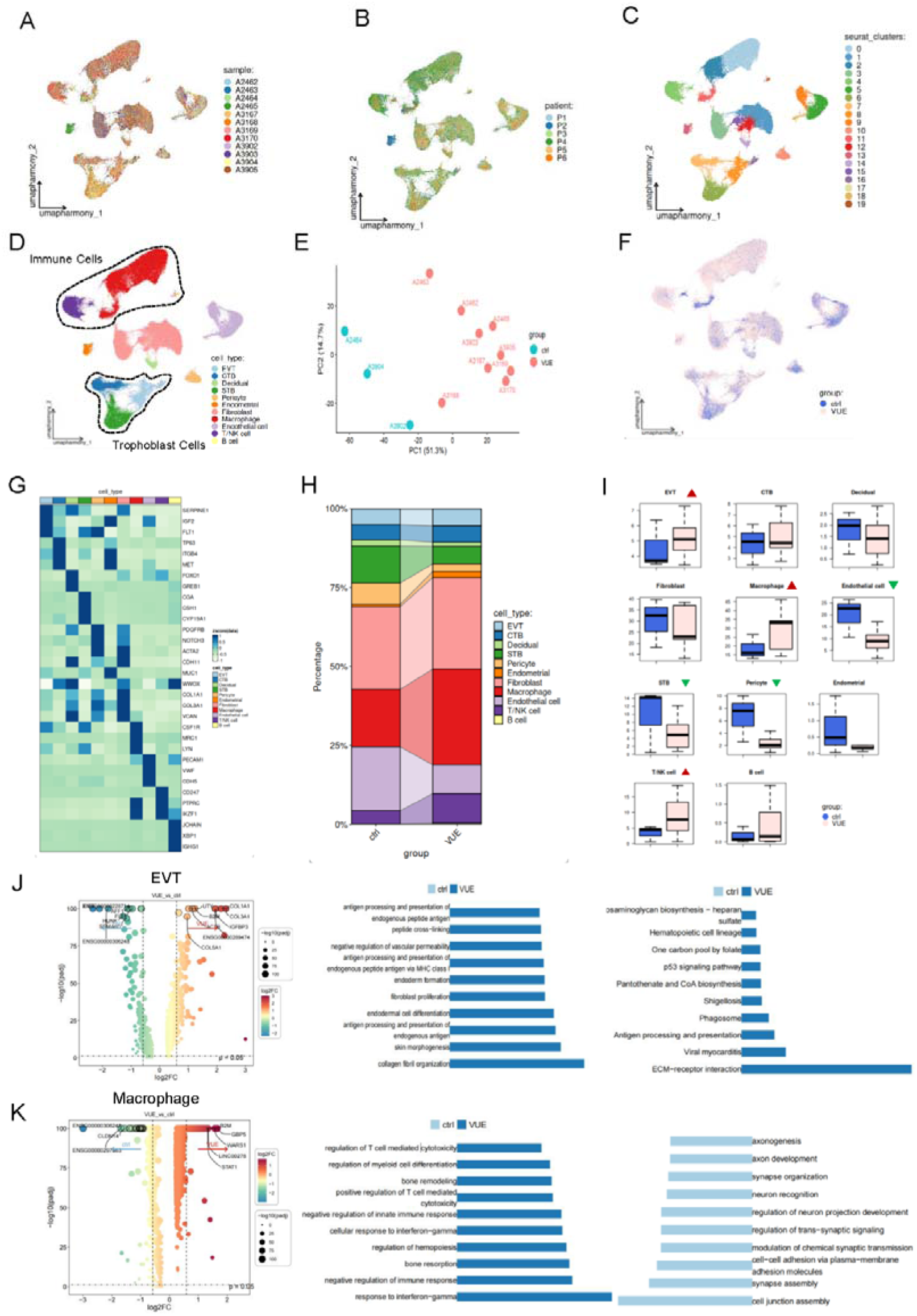
Single-cell transcriptomic landscape in Villitis of Unknown Etiology. (A-C) Uniform Manifold of integrated single-cell data, colored by (A) individual sample ID, (B) patient origin, and (C) unsupervised clusters. (D) Annotated UMAP plot displaying 11 major cell types. (E) Principal Component Analysis (PCA). (F) UMAP visualization colored by group (Control vs. VUE). (G) Heatmap of marker genes. (H) Stacked bar plot of relative cellular composition of each cell subset. (I) Box plots of the cell subset proportions between Control and VUE. Red arrows indicate expanded populations, while green arrows indicate depleted populations. (J-K) Differential expression and functional analysis of EVT and Macrophage. Left: Volcano plot of differentially expressed genes in VUE vs. Control. Right: Bar plot of GO pathway enrichment analysis.

### VUE is associated with global transcriptomic remodeling and selective expansion of immune and extravillous trophoblast compartments

To assess whether disease recurrence influenced placental cellular composition, we compared samples from sporadic and recurrent VUE cases. No significant differences were observed in overall cell-type proportions or global transcriptional patterns between these two groups. Accordingly, all VUE samples were pooled for downstream analyses to maximize statistical power.

Pseudo-bulk principal component analysis revealed clear segregation between control and VUE placentas (Fig. 2E), indicating that VUE is associated with widespread transcriptional remodeling across the placental microenvironment. Consistent with histopathological features of inflammation, VUE placentas exhibited marked enrichment of immune populations, including macrophages, T/NK cells, and B cells (Fig. 2H–2I).

Notably, the extravillous trophoblast (EVT) compartment was also significantly expanded in VUE placentas, whereas stromal and vascular cell populations remained largely unchanged. In contrast, the overall proportion of syncytiotrophoblasts (STBs) was reduced, consistent with villous injury. These findings suggest that VUE is characterized not only by immune infiltration but also by selective remodeling of specific trophoblast lineages.

### VUE-associated transcriptional changes are concentrated in extravillous trophoblasts and macrophages

To identify the principal cellular drivers of VUE-associated transcriptional remodeling, we performed differential expression analysis within individual cell types. Among all placental cell populations, extravillous trophoblasts and macrophages exhibited the most pronounced disease-associated transcriptional changes.

In VUE placentas, EVTs showed significant upregulation of genes involved in antigen processing and presentation, as revealed by Gene Ontology enrichment analysis (Fig. 2J). Under physiological conditions, EVTs exhibit restricted antigen-presenting capacity to maintain maternal–fetal immune tolerance. Activation of these pathways therefore indicates a deviation from the immune-privileged trophoblast state.

In parallel, macrophages displayed robust enrichment of inflammatory and innate immune signaling pathways, including cytokine-mediated responses and leukocyte activation (Fig. 2K). The convergence of antigen presentation programs in EVTs and inflammatory activation in macrophages suggests coordinated trophoblast–immune remodeling as a defining feature of VUE.

### Aberrant MHC class I expression identifies immunogenic trophoblast subpopulations in VUE

Guided by the enrichment of antigen presentation pathways, we examined the expression of major histocompatibility complex (MHC) genes across trophoblast populations. Compared with controls, VUE placentas exhibited significant upregulation of classical MHC class I genes (HLA-A, HLA-B, and HLA-C), together with increased expression of MHC class II–associated genes (HLA-DRA, HLA-DPA1, and CIITA) (Fig. 3A).

**Figure 3.**
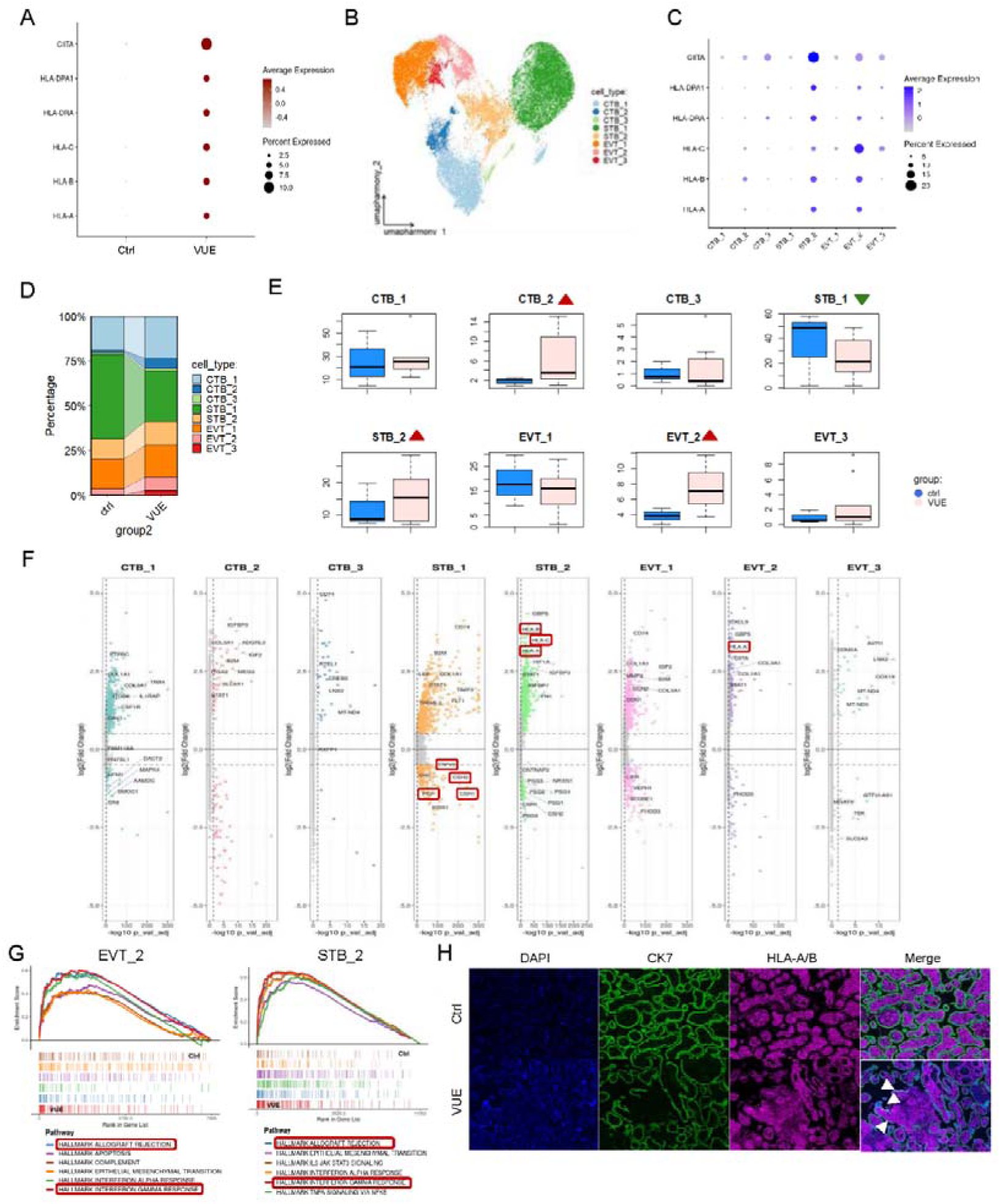
Landscape of trophoblast cells in VUE. (A) Expression of MHC class I and II molecules in VUE trophoblasts compared to controls. (B) UMAP visualization of trophoblast re-clustering, identifying 8 distinct subpopulations. (C) Bubble chart of MHC expression across subclusters. (D) Stacked bar plot of relative cellular composition of each trophoblasts cell subset. (E) Box plots of the trophoblasts cell subset proportions between Control and VUE. Red arrows indicate expanded populations, while green arrows indicate depleted populations. (F) Volcano plots of trophoblasts cell subset differentially expressed genes. (G) Gene Set Enrichment analysis of EVT_2 and STB_2. (H) Multiplex immunohistochemistry (mIHC) showing CK7^+^ (green) HLA-A/B^+^ (magenta) trophoblasts in VUE tissue (white arrows).

To resolve trophoblast heterogeneity, we re-clustered trophoblast cells and identified eight transcriptionally distinct subpopulations, comprising three cytotrophoblast (CTB), two syncytiotrophoblast (STB), and three extravillous trophoblast (EVT) subsets (*17*)(Fig. 3B). Mapping MHC gene expression across these subsets revealed that classical MHC class I expression was largely confined to specific subpopulations, most prominently EVT_2 and STB_2 (Fig. 3C).

Quantitative analysis demonstrated that the overall expansion of EVTs in VUE placentas was driven primarily by selective enrichment of the MHC-high EVT_2 subset (Fig. 3D–3E). Although the total STB compartment was reduced in VUE, the relative proportion of the MHC-high STB_2 subset within the remaining STBs was increased, indicating preferential persistence of immunogenic syncytiotrophoblasts.

### Functional impairment of syncytiotrophoblasts and inflammatory activation of MHC-high trophoblast subsets

To assess the functional consequences of STB remodeling, we examined gene expression changes within the MHC-low STB_1 subset. In VUE placentas, STB_1 cells exhibited marked downregulation of genes critical for placental endocrine and transport functions, including *CSH1*, *CSH2*, *PGF*, and *TRPV6* (Fig. 3F), indicating functional compromise of the remaining syncytiotrophoblast layer.

In contrast, gene set enrichment analysis of the MHC-high EVT_2 and STB_2 subsets revealed significant enrichment of interferon-γ response and allograft rejection pathways (Fig. 3G). These transcriptional programs are characteristic of immune activation and are inconsistent with the maintenance of trophoblast immune quiescence. Multiplex immunohistochemical analysis confirmed the presence of CK7CHLA-A/BC trophoblasts within VUE lesions, providing spatial validation of aberrant antigen-presenting trophoblast subsets in situ (Fig. 3H).

### Maternal macrophage infiltration and inflammatory reprogramming of fetal Hofbauer cells in VUE

To further characterize immune remodeling in VUE placentas, we performed sub-clustering of immune populations. This analysis identified macrophages, fetal Hofbauer cells (HBCs), T cells, regulatory T cells, dendritic cells, B cells, and natural killer (NK) cells (Fig. 4A).

**Figure 4.**
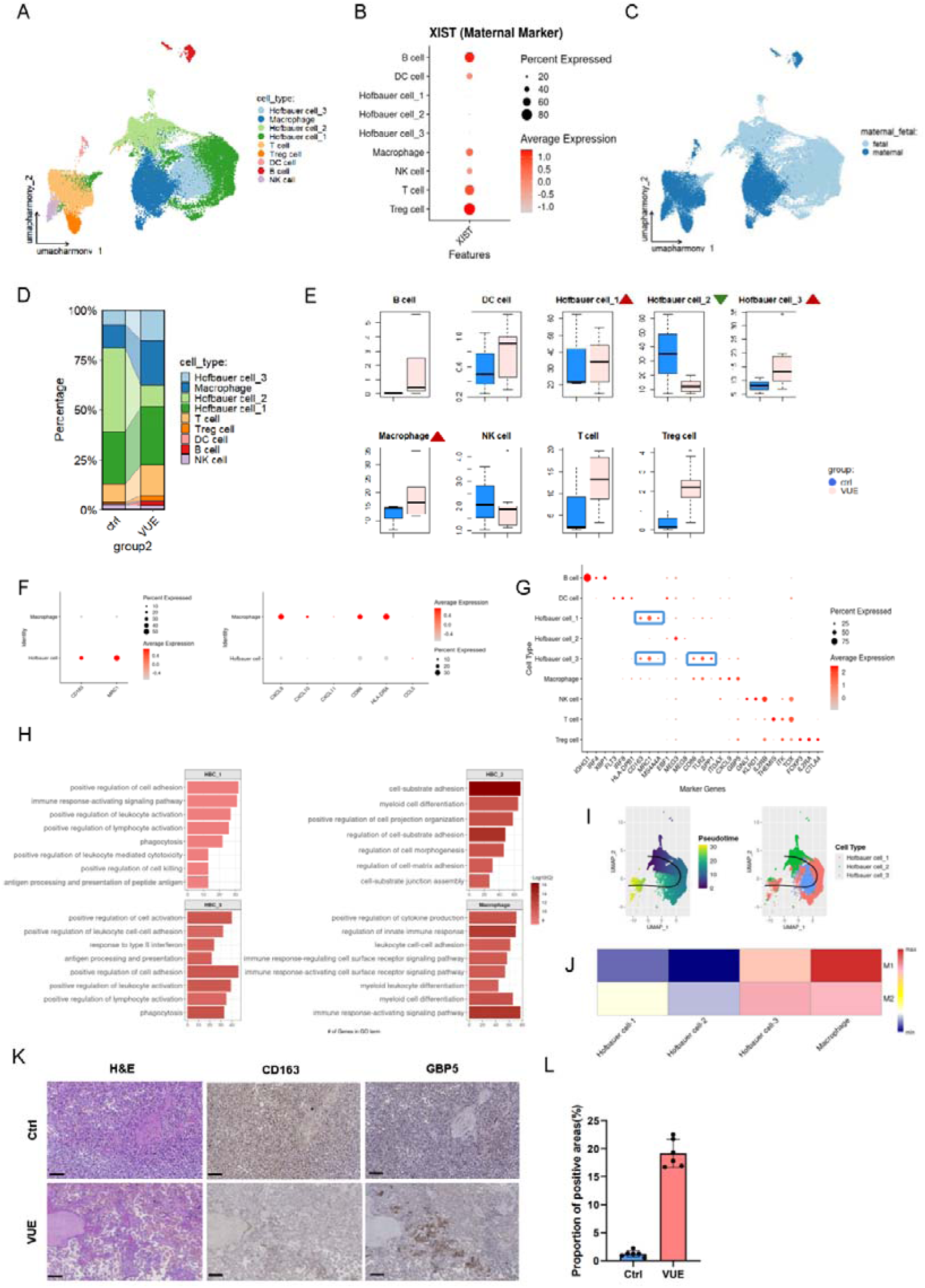
Landscape of immune cells in VUE. (A) UMAP visualization of the immune cells re-clustering, identifying 9 distinct subpopulations. (B) Bubble chart of *XIST* expression in male fetus. (C) UMAP colored by origin. (D) Stacked bar plot of relative cellular composition of each immune cell subset. (E) Box plots of the trophoblasts cell subset proportions between Control and VUE. Red arrows indicate expanded populations, while green arrows indicate depleted populations. (F-G) Phenotypic characterization using polarization markers (F) and cell type markers (G). (H) GO pathway enrichment analysis of HBC and Macrophage. (I) Pseudotime trajectory analysis of HBC. (J) Heatmaps showing M1/M2 polarization of macrophage subtypes calculated by GSVA. (K) Representative IHC images of CD163 (M2 marker) and GBP5 (M1 marker). (L) Quantification confirms the significant increase of GBP5+ inflammatory areas in VUE.

Using XIST expression to distinguish maternal and fetal origin, we confirmed that HBCs were fetal-derived, whereas the majority of infiltrating macrophages were maternal-derived (Fig. 4B–4C). Quantitative analysis revealed a significant increase in maternal macrophages in VUE placentas (Fig. 4D–4E). These cells exhibited a pronounced pro-inflammatory phenotype, characterized by expression of canonical M1-associated markers and enrichment of inflammatory pathways (Fig. 4G, 4J).

In contrast, the overall proportion of fetal HBCs was reduced in VUE. Re-clustering identified three HBC subsets: an M2-like population expressing CD163 and MRC1 (HBC_1), a quiescent homeostatic subset lacking overt inflammatory features (HBC_2), and an activated subset co-expressing inflammatory and immune regulatory genes (HBC_3) (Fig. 4F–4H). VUE placentas showed selective depletion of the quiescent HBC_2 subset and relative enrichment of the activated HBC_3 subset (Fig. 4E).

Pseudotime trajectory analysis revealed a differentiation continuum from HBC_2 toward HBC_3 (Fig. 4I), indicating inflammatory reprogramming of fetal macrophages rather than selective replacement. Immunohistochemical validation further supported this shift, showing reduced CD163C areas and increased GBP5C inflammatory regions in VUE placentas (Fig. 4K–4L).

### Aberrant EVT**–**NK cell interactions define a cytotoxic immune recognition axis in VUE

To determine how immunogenic trophoblasts interact with maternal immune cells, we inferred cell–cell communication networks using CellphoneDB. Global analysis revealed a marked increase in both the number and strength of intercellular interactions in VUE compared with controls (Fig. 5A). Notably, trophoblast populations exhibited increased outgoing and incoming signaling strength, indicating their active participation in the inflammatory microenvironment (Fig. 5B).

**Figure 5.**
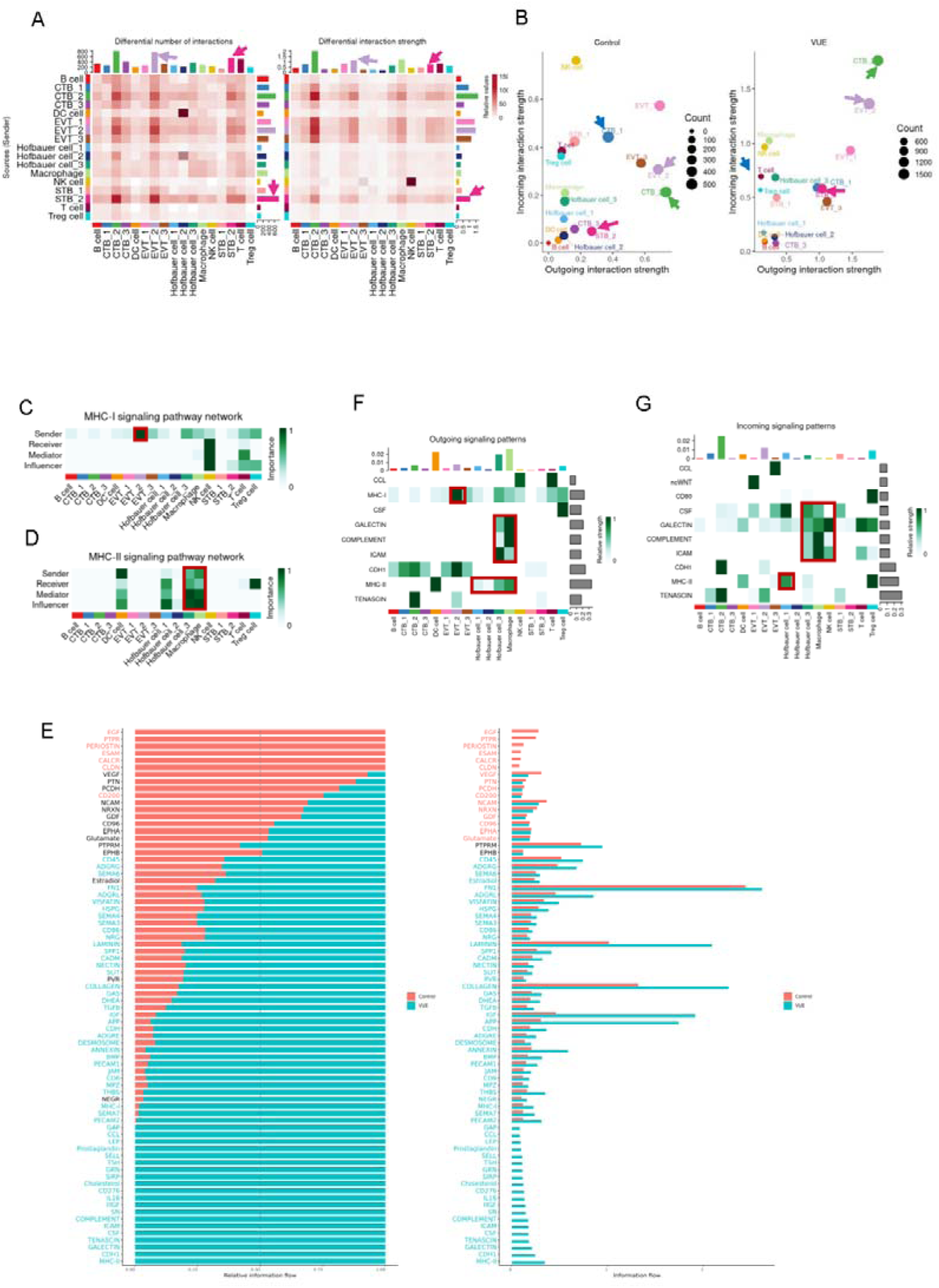
Cell communication between trophoblast cells and immune cells. (A) Heatmaps showing the differential number (left) and interaction strength (right) of signaling pairs between VUE and Control. (B) Scatter plots comparing incoming and outgoing signaling strength in Control and VUE. (C-D) Signaling network visualization of the (C) MHC-I and (D) MHC-II pathways. (E) Bar plots ranking signaling pathways based on relative information flow. Red bars indicate pathways significantly enriched in VUE, while blue bars indicate pathways enriched in Control. (F-G) Pattern recognition analysis identifying dominant outgoing and incoming signaling patterns.

Focusing on MHC class I–mediated signaling, EVT_2 cells emerged as the dominant signal-sending population within the MHC-I network (Fig. 5C). Strikingly, NK cells were identified as the principal signal-receiving population, establishing a specific EVT_2–NK interaction axis in VUE placentas. This interaction was minimal or absent in control placentas, consistent with preservation of trophoblast immune privilege under physiological conditions.

### Immune**–**immune communication networks sustain chronic inflammation in the VUE microenvironment

In contrast to trophoblast-centered MHC-I signaling, MHC class II–mediated interactions were largely restricted to immune–immune networks. Activated fetal HBC_3 cells and maternal macrophages served as the dominant senders and receivers within the MHC-II signaling network (Fig. 5D), indicating reciprocal interactions that reinforce immune activation.

Pathway-level analysis identified a discrete set of signaling pathways selectively enriched in VUE, which could be grouped into four functional modules: immune recruitment and chemotaxis (including CCL and complement signaling), antigen presentation and co-stimulation (MHC-II and CD80), adhesion and extracellular matrix remodeling (ICAM, cadherin, galectin, and tenascin), and myeloid regulation and metabolic support (CSF, GRN, and cholesterol-related pathways) (Fig. 5E).

Outgoing inflammatory and chemotactic signals were primarily driven by maternal macrophages and HBC_3 cells, whereas incoming signals were preferentially received by these same myeloid populations and by NK cells, indicating self-reinforcing inflammatory circuits (Fig. 5F–5G). In contrast, the quiescent HBC_2 subset exhibited minimal participation in these networks.

### Limited viral transcript enrichment in trophoblasts is associated with stress-response signaling in VUE

Although VUE is widely regarded as an immune-mediated placental disorder, infectious or stress-related triggers have been proposed as potential upstream contributors. To explore this possibility in a hypothesis-generating manner, we screened our snRNA-seq data for transcripts derived from a panel of Herpesviridae viruses previously implicated in reproductive pathology.

Viral transcripts were detected at low abundance and were predominantly localized to trophoblast populations, particularly EVTs and STBs (Fig. 6A–6C). While the overall proportion of virus-positive cells was modest, VUE placentas exhibited a relative increase in viral transcript detection within these trophoblast subsets compared with controls. Importantly, total cellular UMI counts were comparable between virus-positive cells from control and VUE samples, whereas viral-specific UMIs were modestly increased in VUE-derived trophoblasts (Fig. 6D).

**Figure 6.**
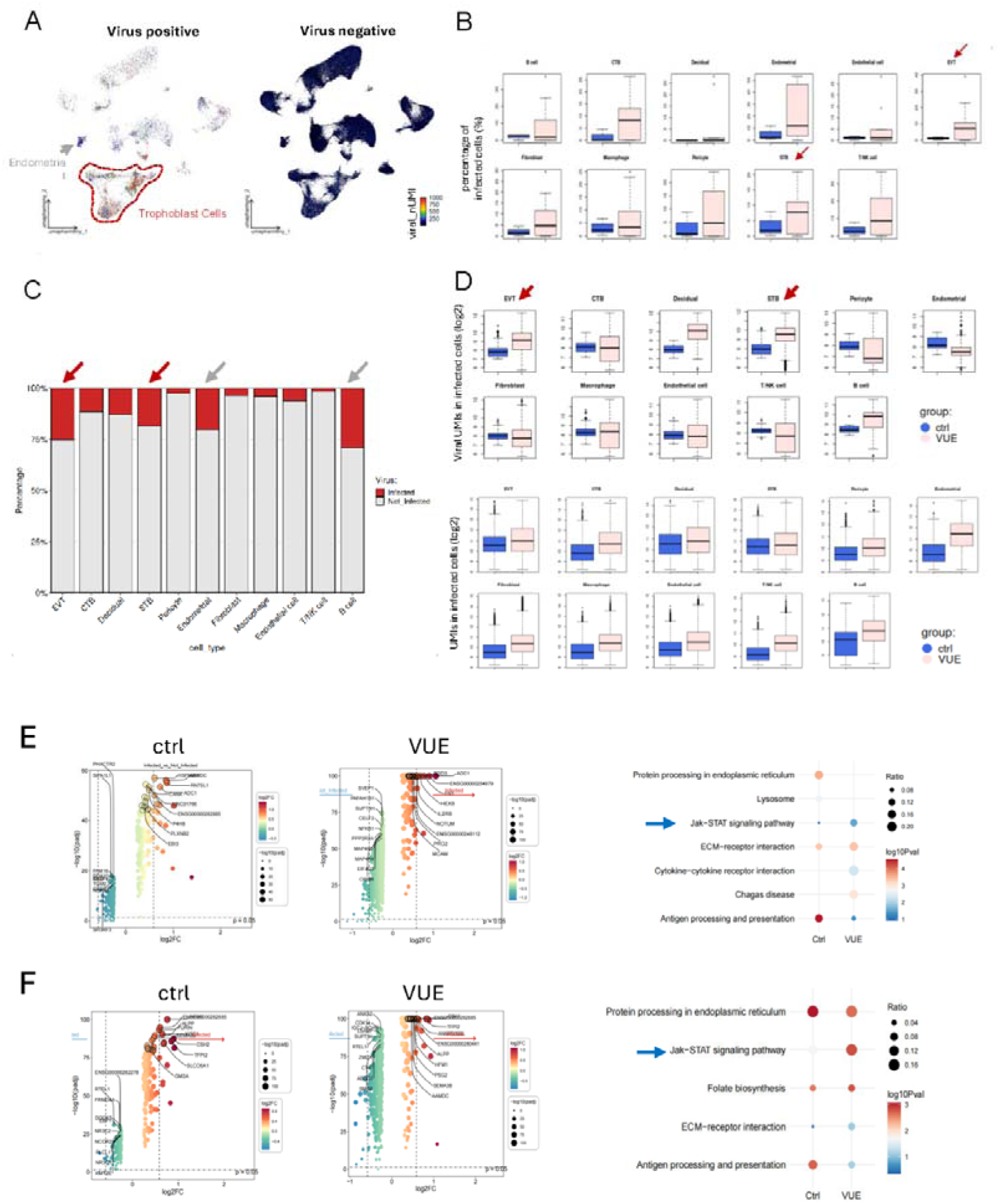
Occult viral infection in VUE. (A) Feature plots mapping viral transcript distribution on the UMAP projection. (B) Box plots demonstrate a proportion of viral-positive cells in VUE compared to controls. (C) Stacked bar plot of relative proportion of viral-positive cells of each immune cell subset. (D) Box plots show UMI and viral-specific UMI counts in VUE. (E-F) Differential expression and functional analysis of EVT and STB. Left: Volcano plot of differentially expressed genes in VUE vs. Control. Right: Bar plot of GO pathway enrichment analysis.

Gene set enrichment analysis revealed activation of JAK–STAT signaling pathways in EVT and STB subsets from VUE placentas (Fig. 6E–6F), consistent with a cellular stress or interferon-associated response. Notably, this transcriptional signature overlapped with that observed in MHC class I–high trophoblast populations. Together, these observations suggest that low-level viral transcription or related intracellular stress responses may be associated with trophoblast immune activation in VUE, without implying a direct infectious etiology.

## DISCUSSION

In this study, we applied an FFPE-compatible single-nucleus RNA sequencing strategy to paired placental samples from sequential pregnancies to systematically dissect the cellular and molecular landscape of villitis of unknown etiology (VUE). By minimizing inter-individual heterogeneity, our analysis reveals a coherent pathogenic framework in which loss of trophoblast immune privilege initiates maternal immune recognition, while coordinated remodeling of maternal and fetal macrophage populations amplifies and sustains chronic placental inflammation. These findings position VUE as an active immune-mediated disorder driven by specific trophoblast–immune interactions, rather than a nonspecific inflammatory response.

Classical histopathology considers the fundamental morphological characteristics of VUE to be the infiltration of immune cells within the villous stroma, accompanied by damage to the villous structure. Severe cases are associated with adverse outcomes such as fetal growth restriction and stillbirth(*18*). Previous immunohistochemical and quantitative studies have further indicated that CD8C T cells and macrophages are enriched in VUE lesions, and that focal placental damage can affect the terminal villi, a critical structure for substance exchange. (14–16) This study observed an increase in the proportion of immune cells at single-cell resolution, accompanied by a decrease in trophoblasts, which is highly consistent with the histological framework of immune infiltration and damage to villous functional units. Simultaneously, this study localized these changes to specific cell lineages and subpopulations, providing more direct evidence to explain the functional consequences of VUE. (11–16)Two major hypotheses have long been proposed to explain the pathogenesis of VUE: one attributing the lesion to persistent or occult infection–driven chronic inflammation(*19*), and the other positing maternal immune rejection of fetal antigens as the dominant mechanism(*6*, *20*). Although early studies favored infectious triggers, accumulating transcriptomic and multiplex immunohistochemical evidence increasingly supports the immune rejection model. VUE lesions consistently exhibit upregulation of the chemokines CXCL9 and CXCL10, enhanced recruitment of T cells, and enrichment of pathways associated with tissue rejection and allogeneic immune responses(*21*, *22*).

Our data provide cell type–resolved mechanistic support for a rejection-like process. Differential gene expression and gene set enrichment analyses highlighted robust activation of antigen processing and presentation pathways together with inflammatory signaling in extravillous trophoblasts (EVTs), while macrophages displayed pronounced upregulation of cytokine production and inflammatory response programs. These findings refine the concept of immune rejection in VUE by defining it as a cooperative process involving increased trophoblast immunogenicity coupled with myeloid-driven amplification of inflammation, rather than a phenomenon driven solely by infiltrating lymphocytes.Mechanistically, both trophoblasts and myeloid cells are known to upregulate antigen presentation machinery in interferon-rich inflammatory environments, thereby increasing the probability of maternal T cell recognition and establishing positive feedback loops of immune activation(*7*, *21*, *23*). Katzman and colleagues reported enhanced STAT1 phosphorylation in trophoblasts within VUE lesions, implicating local IFN/STAT1 signaling in lesion formation(*24*). This signaling axis is directionally consistent with the JAK–STAT and interferon-response pathway enrichment observed in our study, supporting a model in which VUE represents a sustained, multi-lineage immune activation state involving trophoblast–myeloid–lymphoid crosstalk.

Successful human pregnancy relies on robust immune tolerance that permits prolonged coexistence of a semi-allogeneic fetus within the maternal host(*25*). This balance is maintained in part by the unique HLA expression repertoire of trophoblasts, particularly EVTs(*26*, *27*), which express HLA-C and the non-classical HLA class Ib molecules HLA-E, -F, and -G that interact with maternal uterine NK cell receptors to coordinate trophoblast invasion, vascular remodeling, and immune regulation(*28*). Disruption of this finely tuned system is increasingly recognized as a contributor to inflammatory placental disease(*29*).

Within this immunological context, our trophoblast re-clustering analysis identified EVT_2 and STB_2 subpopulations characterized by elevated MHC expression, particularly MHC class I, which were selectively enriched or showed increasing trends in VUE placentas. These findings suggest that a subset of trophoblasts may shift from a low-immunogenic, immunomodulatory state toward a more antigen-presenting, immune-visible phenotype. This interpretation aligns closely with observations by Enninga and colleagues and others demonstrating enhanced HLA class I expression and upregulation of rejection-associated transcriptional programs in VUE placentas(*7*).

The syncytiotrophoblast (STB) layer is the principal mediator of maternofetal exchange and placental endocrine function; its structural and molecular integrity directly determines oxygen and nutrient transport efficiency as well as hormone secretion profiles(*30*, *31*). VUE frequently involves terminal villi and is clinically associated with fetal growth restriction, implicating STB dysfunction as a key downstream consequence(*2*, *32*). In our data, the STB_1 subpopulation exhibited downregulation of genes related to metabolism, endocrine activity, and transport, while an MHC-high STB_2 subpopulation emerged concurrently, suggesting that STBs in VUE may undergo parallel processes of functional decline and inflammatory reprogramming.

Hofbauer cells (HBCs) are fetal-derived, tissue-resident macrophages present early in placental development and involved in angiogenesis, tissue homeostasis, and pathogen defense(*33*, *34*). Recent work has highlighted the coexistence of placenta-associated maternal macrophages with fetal HBCs at the maternal–fetal interface(*35*). Using XIST expression and transcriptional signatures(*28*), we distinguished maternal from fetal macrophages and observed increased infiltration of maternal macrophages together with shifts in HBC subpopulation composition and pseudotime trajectories, consistent with classical descriptions of maternal immune cell entry into fetal tissues in VUE.

The maternal–fetal interface is inherently a highly interactive cellular network(*28*), relying on tightly regulated ligand–receptor signaling among trophoblasts, NK cells, macrophages, and T cells to maintain homeostasis and vascular remodeling(*36*, *37*). Using CellPhoneDB, we observed a global increase in inferred cell–cell communication in VUE, particularly involving MHC class I and II–related interactions, and identified a putative EVT–NK recognition axis. While ligand–receptor inference reflects interaction potential rather than confirmed signaling events, these analyses elevate VUE from a focal pathological lesion to a disorder of dysregulated multicellular immune networks.

Whether VUE is initiated by occult infection remains unresolved. Systematic evaluations suggest that direct viral detection is inconsistent and unlikely to account for most cases(*19*). However, the maternal–fetal interface mounts strong interferon-mediated innate immune responses to pathogens, and such responses may themselves induce bystander tissue damage and barrier dysfunction(*38*, *39*). In our study, low-level viral transcripts were detected together with activation of JAK–STAT and interferon pathways, a pattern more consistent with an antiviral-like immune activation state than with overt infection.

Several limitations should be acknowledged. Although the paired-sample design strengthens mechanistic inference, the overall cohort size remains modest, limiting the resolution of rare cell populations and inter-individual variability. RNA degradation inherent to FFPE specimens may bias detection toward more abundant transcripts, and the absence of spatially resolved transcriptomic data constrains direct mapping of molecular changes to histological lesions. Future studies integrating spatial transcriptomics, proteomics, and functional assays will be essential to further validate and extend these findings.

In summary, our study defines VUE as a disorder of disrupted maternal–fetal immune tolerance driven by aberrant trophoblast antigen presentation and sustained by macrophage-mediated inflammatory amplification. By establishing a mechanistic link between trophoblast differentiation, immune recognition, and placental injury, these findings advance our understanding of VUE pathogenesis and provide a conceptual framework for future diagnostic and therapeutic exploration.

## MATERIALS AND METHODS

### Ethics approval

This study protocol about human tissue specimens was approved by the Ethics Committee of Jinan Maternity and Child Care Hospital Affiliated to Shandong First Medical University (No. KY R-24-134). All patients provided written informed consent.

### Patients and Samples

12 FFPE placenta samples from 6 females were retrospectively collected at Jinan Maternity and Child Care Hospital Affiliated to Shandong First Medical University. Each of the female had two pregnancies. Three of them had no VUE at the first pregnancy, while have VUE at the second pregnancy. The other three had VUE at both pregnancies. Villitis of unknown etiology (VUE) was diagnosed and graded according to the Amsterdam criteri. High-grade VUE was defined as inflammation involving ≥10 contiguous villi in at least one focus or as patchy/diffuse inflammation affecting multiple separate foci, whereas low-grade VUE was defined as inflammation involving <10 contiguous villi per focus, with more than one focus required for diagnosis. All cases were high-level VUE.

### Histology and Immunohistochemical staining

The placenta tissue embedded in paraffin, cut into 4 μm sections, and dewaxed with xylene. Hematoxylin/eosin (H&E) staining of the sections was performed according to standard staining procedures. For immunohistochemistry, primary antibodies against human GBP5 (13220-1-AP, Proteintech), and CD163 (16646-1-AP, Proteintech) were used. For multiplex immunohistochemical staining, primary antibodies are against human CK7 (17513, Proteintech), and HLA-A/B (15240-1-AP, Proteintech), the Four Detect Kit for Rabbit Primary Antibody kit (PK10033, Proteintech) used for multiple staining. All the sections were scanned via a KF-PRO-005 Digital Slide Scanner (KFBIO), and ImageJ was used to calculate the area ratio of positive areas in the sections.

### scRNA-seq Data Analysis

#### Quality Control and Preprocessing

Single-cell RNA sequencing (scRNA-seq) data analysis was performed in R (version 4.4.1) using the Seurat package (version 5.0.0). Raw sequencing data from 12 samples (3 control and 9 VUE cases) were processed using the Cell Ranger pipeline. For each sample, gene expression matrices were imported using the Read10X function and Seurat objects were created with initial filtering parameters (min.cells = 3, min.features = 200). Quality control metrics were computed for each cell, including the percentage of mitochondrial reads using the PercentageFeatureSet function with the pattern "^MT-". Cells were retained if they expressed between 200 and 6,000 genes (nFeature_RNA) and had less than 20% mitochondrial read mapping. Genes expressed in fewer than three cells were excluded from downstream analyses.

### Normalization and Integration

Normalization was performed using the SCTransform workflow with the glmGamPoi method, regressing out the percentage of mitochondrial reads to mitigate technical variation. Principal component analysis (PCA) was performed on the normalized data, and the optimal number of principal components was determined using elbow plots. Initial concatenation of cells across the 12 samples revealed residual batch effects. To address this, batch effect correction was performed using the Harmony package (version 1.2.1) with the following parameters: theta = 2, lambda = 0.5, and max_iter = 25. The integration was performed on sample identity (orig.ident) to correct for inter-sample technical variation while preserving biological differences between conditions.

### Dimensionality Reduction and Clustering

Uniform Manifold Approximation and Projection (UMAP) was performed on the Harmony-corrected embeddings using the first 30 principal components for visualization. Unsupervised clustering was performed using the FindNeighbors function (30 principal components) followed by the FindClusters function with a resolution of 0.5, which identified 20 distinct clusters (clusters 0-19). Cluster-specific marker genes were identified using the FindAllMarkers function with the MAST statistical framework, requiring genes to be expressed in at least 25% of cells within a cluster (min.pct = 0.25) and have a minimum log2 fold change of 0.5 (logfc.threshold = 0.5). Cell type identities were manually assigned based on canonical marker genes derived from previous placental single-cell studies(*17*), resulting in 12 major cell types: Hofbauer cells, macrophages, T/NK cells, B cells, fibroblasts, endothelial cells, pericytes, decidual cells, endometrial cells, cytotrophoblasts (CTB), syncytiotrophoblasts (STB), and extravillous trophoblasts (EVT).

### Pseudobulk Analysis and Sample-Level Visualization

To visualize sample-level transcriptional differences between VUE and control groups, pseudobulk expression profiles were generated by aggregating raw counts across all cells within each biological sample. Variance-stabilizing transformation (VST) was applied using DESeq2 (version 1.44.0) to normalize the pseudobulk expression matrix. To account for potential confounding factors, surrogate variable analysis (SVA) was performed using the sva package, identifying and removing three surrogate variables using the removeBatchEffect function from limma. PCA was then performed on the batch-corrected data to visualize sample separation between VUE (n=9) and control (n=3) groups.

### Immune Cell Subset Analysis

Immune cells (Hofbauer cells, macrophages, T/NK cells, and B cells) were extracted from the integrated dataset for detailed subset analysis. The immune submatrix underwent re-normalization using SCTransform followed by batch effect correction with Harmony using identical parameters as the main analysis (theta = 2, lambda = 0.5, max_iter = 25). UMAP visualization was generated using the first 30 Harmony-corrected components. Unsupervised clustering at resolution 0.4 identified 15 clusters, which were manually annotated into nine immune cell subtypes based on established marker gene expression: three Hofbauer cell subtypes (Hofbauer cell_1, Hofbauer cell_2, Hofbauer cell_3), monocyte-derived macrophages, T cells, regulatory T cells (Treg), NK cells, B cells, and dendritic cells (DC).

The maternal versus fetal origin of immune cells was determined using XIST expression analysis. Samples from male fetuses were subset to assess XIST expression patterns across immune cell populations. Hofbauer cells (all three subtypes) were classified as fetal-derived, while macrophages, T cells, Treg cells, NK cells, B cells, and DC cells were classified as maternal-derived based on XIST expression patterns. Cell type proportions were calculated for each sample and compared between VUE and control groups. Gene Ontology (GO) enrichment analysis was performed on cluster-specific marker genes for each Hofbauer cell subtype and macrophage population using the clusterProfiler R package (version 4.2.2) to characterize their functional states.

### Macrophage Polarization Analysis

To assess macrophage polarization states, Gene Set Variation Analysis (GSVA) was performed using the GSVA package with predefined M1 and M2 gene signatures. The M1 signature included: IL23, TNF, CXCL9, CXCL10, CXCL11, CD86, IL1A, IL1B, IL6, CCL5, IRF5, IRF1, CD40, IDO1, KYNU, and CCR7. The M2 signature included: IL4R, CCL4, CCL13, CCL20, CCL17, CCL18, CCL22, CCL24, LYVE1, VEGFA, VEGFB, VEGFC, VEGFD, EGF, CTSA, CTSB, CTSC, CTSD, TGFB1, TGFB2, TGFB3, MMP14, MMP19, MMP9, CLEC7A, WNT7B, FASL, TNFSF12, TNFSF8, CD276, VTCN1, MSR1, FN1, and IRF4. GSVA enrichment scores were computed using the gsvaParam function on averaged expression profiles of each macrophage subtype (Hofbauer cell_1, Hofbauer cell_2, Hofbauer cell_3, and macrophages) and visualized as a heatmap.

### Trajectory Inference

Hofbauer cell developmental trajectories were reconstructed using the Slingshot R package (version 2.12.0). The analysis utilized the expression matrix of three Hofbauer cell subtypes with unmerged cluster labels and UMAP embeddings as input. Pseudotime values were computed along inferred lineages to characterize the developmental progression of Hofbauer cells.

### Trophoblast Cell Subset Analysis

Trophoblast cells (CTB, STB, and EVT) were extracted for detailed subset analysis. Re-normalization and batch correction were performed as described above. UMAP visualization was generated using 30 Harmony-corrected components. Unsupervised clustering at resolution 0.3 identified 10 clusters, which were manually annotated into eight trophoblast subtypes: three CTB subtypes (CTB_1, CTB_2, CTB_3), two STB subtypes (STB_1, STB_2), and three EVT subtypes (EVT_1, EVT_2, EVT_3) based on marker gene expression. Expression of MHC class I (HLA-A, HLA-B, HLA-C), MHC class II (HLA-DRA, HLA-DPA1), and CIITA genes was visualized using dot plots to assess antigen presentation capacity across conditions and trophoblast subtypes. Cell type proportions were compared between VUE and control groups.

### Differential Expression Analysis

Differential gene expression analysis between VUE and control groups was performed for each cell type using the MAST algorithm implemented in the FindMarkers function of Seurat. Genes with false discovery rate (FDR)-adjusted p-values < 0.05 were considered differentially expressed. Results were visualized using volcano plots with -log10(adjusted p-value) on the x-axis and log2 fold change on the y-axis for each trophoblast subtype. Key differentially expressed genes were highlighted, including genes involved in immune response (HLA-A, HLA-B, HLA-C, STAT1, CXCL9), extracellular matrix organization (COL1A1, COL3A1), and placental hormones (CSH1, CSH2, PSG family genes). Gene Ontology (GO) enrichment analysis of differentially expressed genes was conducted using the clusterProfiler R package (version 4.2.2).

### Gene Set Enrichment Analysis

Gene Set Enrichment Analysis (GSEA) was performed to identify Hallmark path ways differentially enriched between VUE and control samples in STB_2 and EV T_2 cell populations, which showed the most pronounced transcriptional changes. Pseudobulk expression matrices were generated by aggregating raw counts across cells within each biological sample for each cell type (VUE: n=9; control: n=3). Differential expression analysis was conducted using DESeq2, with genes filtered to require at least 10 counts in at least 3 samples. Genes were ranked by Wald statistics from the DESeq2 analysis. GSEA was implemented using the fgsea R pac kage (version 1.30.0) with Hallmark gene sets from MSigDB (msigdbr package), employing 10,000 permutations, minimum gene set size of 15, and maximum gen e set size of 500. Pathways with FDR < 0.05 were considered significantly enriched. Enrichment plots were generated showing running enrichment scores for selected pathways including HALLMARK_INTERFERON_GAMMA_RESPONSE, HA LLMARK_ALLOGRAFT_REJECTION, HALLMARK_INTERFERON_ALPHA_R ESPONSE, HALLMARK_APOPTOSIS, HALLMARK_EPITHELIAL_MESENCH YMAL_TRANSITION, and HALLMARK_COMPLEMENT.

### Cell-Cell Communication Analysis

Cell-cell communication networks were inferred using the CellChat R package (version 2.2.0) to characterize ligand-receptor interactions between trophoblast and immune cell populations. Integrated trophoblast and immune cell datasets were separated by condition (VUE and control), and CellChat objects were created using the human ligand-receptor interaction database (CellChatDB.human). For each condition, overexpressed ligands and receptors were identified, and communication probabilities were computed using the truncated mean method (type = "truncatedMean", trim = 0.1) for robust estimation. Interactions involving fewer than 10 cells were filtered out (min.cells = 10). Communication networks were aggregated at the pathway level to reveal intercellular communication patterns.

For comparative analysis between conditions, CellChat objects were merged after lifting the control dataset to match cell type labels in the VUE dataset using the liftCellChat function. Differential analysis included quantification of interaction numbers and strengths, visualized as heatmaps showing the differential number of interactions and differential interaction strength between conditions. Signaling roles of each cell type were assessed by computing network centrality measures and visualized using scatter plots comparing outgoing and incoming interaction strengths. Information flow analysis was performed using the rankNet function to identify signaling pathways with significant changes between conditions. Signaling role networks for specific pathways (MHC-I and MHC-II) were generated using the netAnalysis_signalingRole_network function to identify sender, receiver, mediator, and influencer cell types. Outgoing and incoming signaling patterns for selected pathways (CCL, MHC-I, CSF, GALECTIN, COMPLEMENT, ICAM, CDH1, MHC-II, TENASCIN) were visualized using heatmaps generated by the netAnalysis_signalingRole_heatmap function.

### Statistical Analysis

All statistical analyses were performed in R (version 4.4.1). For differential expression analysis, the MAST framework was used with FDR correction for multiple testing (Benjamini-Hochberg method). For GSEA, statistical significance was assessed using 10,000 permutations with FDR < 0.05 considered significant.

### Software and Packages

Key R packages used in this study include: Seurat (v5.0.0) for single-cell data processing and analysis; Harmony (v1.2.1) for batch effect correction; DESeq2 (v1.44.0) for pseudobulk differential expression analysis; GSVA for gene set variation analysis; fgsea (v1.30.0) for gene set enrichment analysis; clusterProfiler (v4.2.2) for functional enrichment analysis; CellChat (v2.2.0) for cell-cell communication analysis; Slingshot (v2.12.0) for trajectory inference; sva for surrogate variable analysis; limma for batch effect removal; and SCP for visualization.

## Acknowledgements

We thank Prof. Xiaoying Fan at Guangzhou Laboratory for suggestions on data analysis, Dr. Ling Dong at M20 Genomics for technical support on sample processing.

## Author contributions

Study design: F.G, Y.W., YC.W., Y.S;

Sample collection and pathology evaluation: J.L., M.W., T.L, Y.W., F.G.;

Methodology: Q.Z., YC.W.;

Single-nucleus RNA-seq experiments: M.W., Q.Z., Y.S., YC.W.;

Data analysis: J.L., M.W., Q.Z. Y.C., G.Y., Y.S.;

Histology and immunohistochemistry: J.L., M.W., Y.W., F.G.

Visualization: Q.Z., Y.C.

Supervision: F.G, Y.W., YC.W., Y.S;

Writing – original draft: M.W., Q.Z., Y.S;

Writing – review & editing: all authors.

## Data availability

All data needed to evaluate the conclusions in the paper are present in the paper and/or the Supplementary Materials. The scRNA-seq data have been deposited to the Genome Sequence Archive for Human (GSA-Human) of China National Center for Bioinformation (CNCB) database under the accession number HRA016786.

## Figures

**Supplementary Figure 1.**
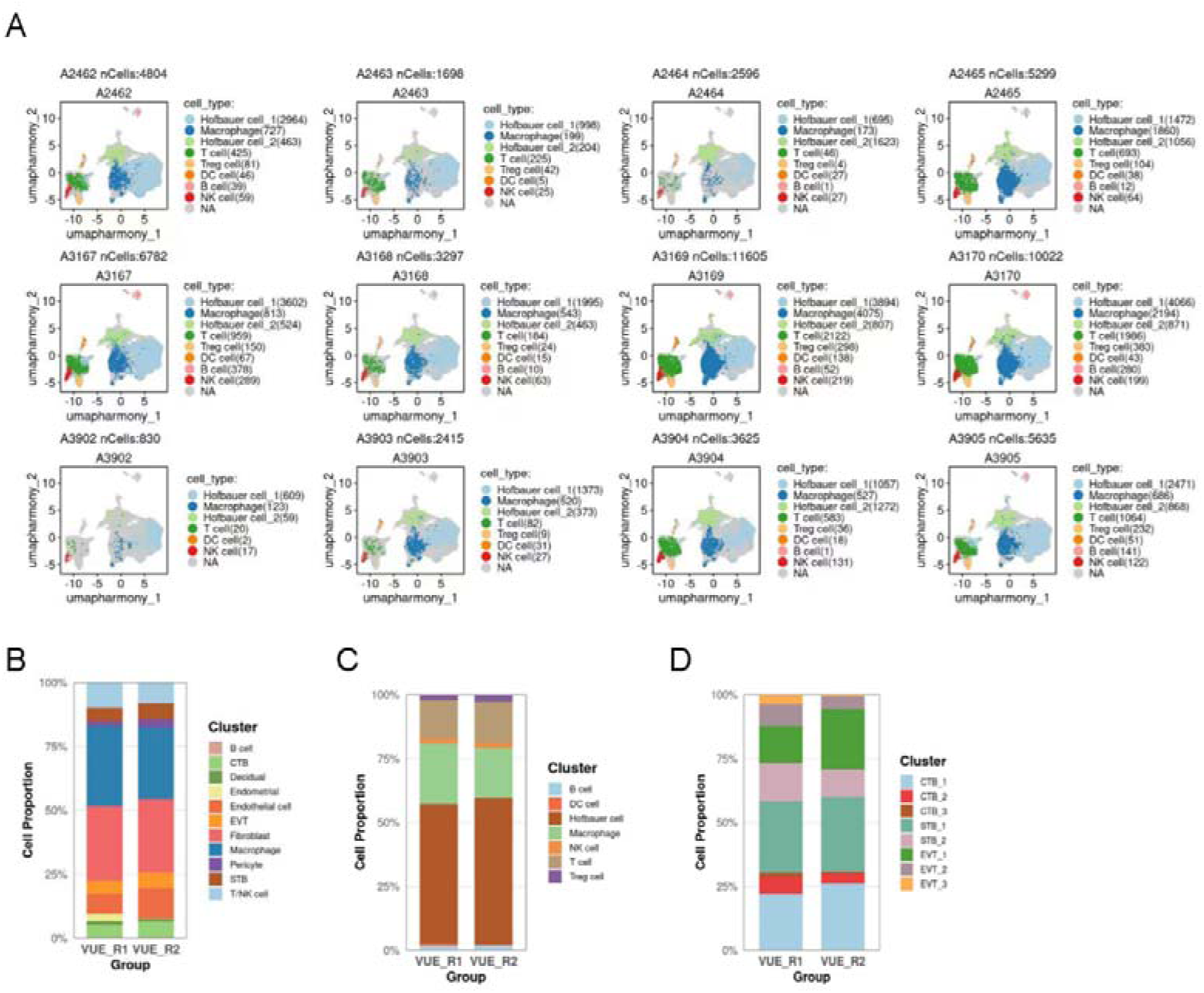
Single-cell landscape and cell type composition analysis across samples. (A) UMAP visualization of cell populations across 12 individuals samples (A2462–A3905). (B-D) Stacked bar plot showing the relative proportions of major cell of recurrent VUE samples. (B) Overall cell proportion composition. (C) Immune cells proportion composition. (D) Trophoblast cells proportion composition.

**Supplementary Figure 2.**
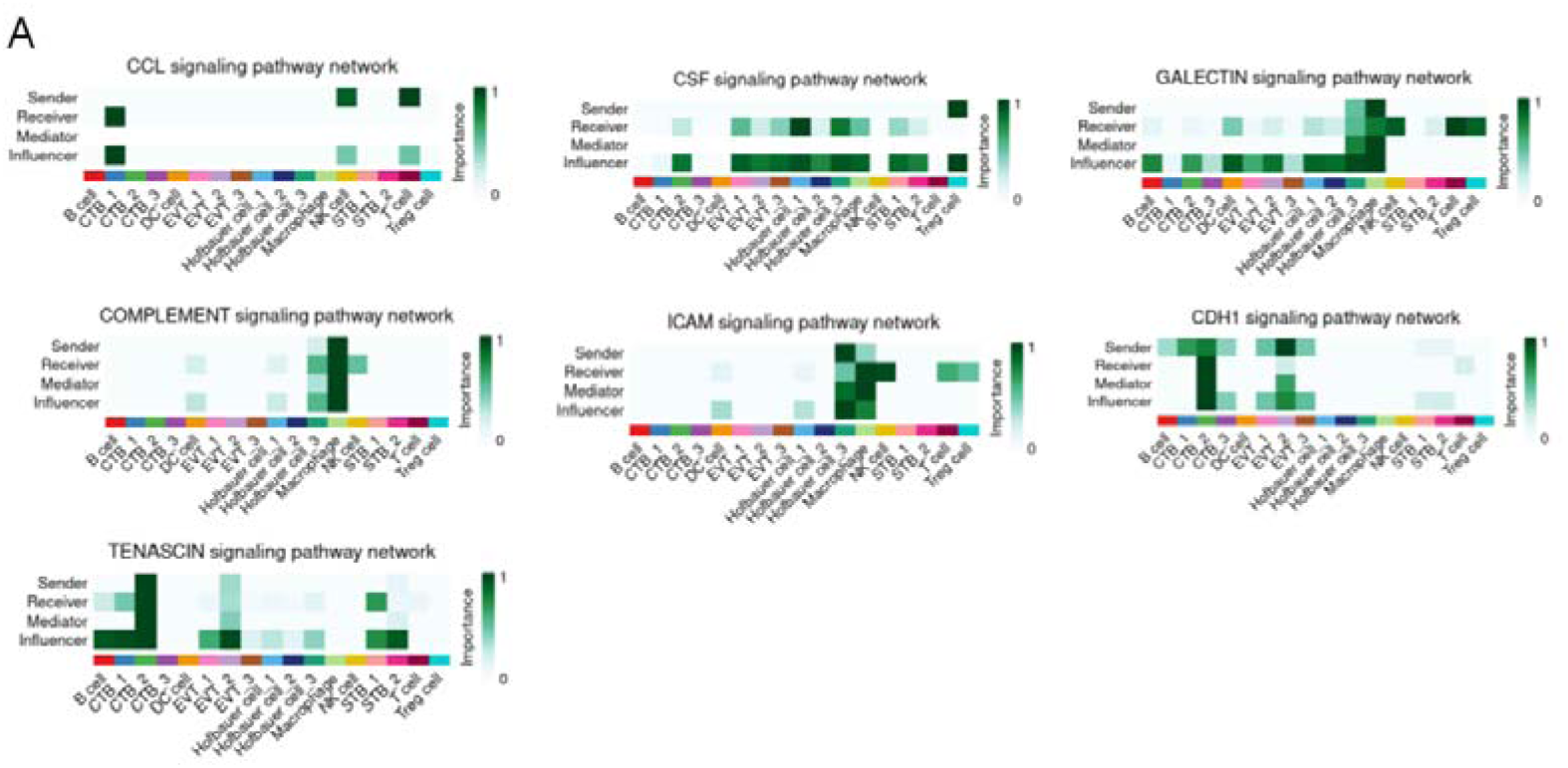
Identification of dominant signaling roles of cell groups in key pathways. **(A)** Heatmaps illustrating the network centrality scores for major signaling pathways (CCL, CSF, GALECTIN, COMPLEMENT, ICAM, CDH1, and TENASCIN).

